# Leaf and cluster spectra reveal trait dependent prediction of grapevine cluster architecture and juice quality

**DOI:** 10.64898/2026.03.27.714894

**Authors:** Carlos A. Robles-Zazueta, Timo Strack, Maximilian Schmidt, Paolo Callipo, Hannah Robinson, Akshaya Vasudevan, Kai P. Voss-Fels

**Affiliations:** Department of Plant Breeding, Hochschule Geisenheim University, 65366 Geisenheim, Germany

**Keywords:** abiotic stress, biotic stress, cluster architecture, grapevine breeding, partial least squares regression, phenomics, spectral reflectance

## Abstract

Grapevine cluster architecture is a key selection target in breeding programs because it influences disease susceptibility, yield stability and juice quality. High-throughput phenotyping offers a rapid and non-destructive approach to capture biochemical and structural variation in these traits, yet the influence of plant organ reflectance and data partitioning strategies on trait prediction remains poorly understood. In this study, we evaluated how hyperspectral reflectance from different grapevine organs contributes to the prediction of cluster architecture and juice quality traits in two clonal populations of Riesling and Pinot. Using partial least squares regression (PLSR), we assessed the prediction accuracy of eight cluster architecture and six juice quality traits under two data partitioning strategies. Models based on cluster reflectance outperformed those using dry leaf reflectance for most traits, except for pH. Partitioning the dataset by cluster type increased trait variance and improved predictions for number of berries (R² = 0.53), berry diameter (R² = 0.79), and total acidity (R² = 0.48). Visible, red-edge and NIR spectra were most informative regions to predict the traits studied. Together, our results highlight the importance of organ-specific data and appropriate calibration strategies to improve phenomic models for the development of scalable proxies for grapevine improvement.

**Highlight:** Spectral phenomics reveals that prediction accuracy in grapevine depends on organ spectral signatures and traits, with cluster reflectance outperforming leaves, informing new phenotyping strategies for breeding improvement.

## 1. Introduction

Increased environmental variability due to climate change is increasing pressure on agricultural systems which are expected to maintain or raise crop productivity with reduced fertilizer and water inputs (Robles-Zazueta et al., 2024). These pressures are exacerbated for perennial crops such as grapevine where cumulative effects of prolonged and increasingly frequent heatwaves, drought periods and inconsistent precipitation patterns across multiple years can reduce photosynthetic performance, shorten phenological stages, alter source-sink dynamics diminishing yield and altering the chemical composition of the berries (Fraga et al., 2016; van Leeuwen et al., 2024).

Higher temperatures and extended heatwaves increase the probability of sunburn damage of the berries (Gambetta et al., 2021). Additionally, alterations in precipitation patterns (e.g. a higher frequency of rain during harvest) create favourable conditions for bunch rot infection caused by *Botrytis cinerea* which may further lead to secondary fungi and bacteria infections (Molitor et al., 2020). These abiotic and biotic pressures affecting grapevine clusters can lead to yield losses and juice quality reduction. This makes the case for characterization of cluster architecture at breeding scale (Herzog et al., 2022).

This challenge is particularly relevant in clonally propagated perennials such as grapevine, where genetic improvement relies on clonal selection and the propagation of somatic variants derived from a single cultivar. Although these clones are genetically similar, phenotypic variations are common due to somatic mutations and epigenetic modifications, which can lead to differences in grapevine agronomic performance and morphological traits, including cluster architecture. Therefore, clonal populations represent a genetically narrow but phenotypically relevant source of variability for important breeding targets (Callipo et al., 2025).

In this context, grapevine cluster architecture is a complex trait that is key in the determination of yield potential, quality and susceptibility to abiotic and biotic stress resistance. This trait is a sum of individual berry traits such as number, diameter, volume and its spatial arrangement (Tello and Ibañez, 2018). Furthermore, it can influence juice quality composition affecting sugar accumulation, phenolic components, organic acids and pH which are paramount traits for winemaking (Zanchin et al., 2025).

Cluster architecture and quality traits are measured destructively using labour and resource intensive protocols limiting its evaluation at large scale and integration into breeding pipelines (Diago et al., 2014). As a result, selection for these traits have relied on visual assessments that can fail to capture the complexity of the relationship between berry, cluster and quality traits (Tello et al., 2015). However, high-throughput phenotyping (HTP) approaches can overcome these limitations with rapid, precise and repeatable plant assessments at organ, canopy or vineyard scales.

In grapevine, HTP efforts have focused on traits related to abiotic stresses, vigour and disease detection using RGB imaging, chlorophyll fluorescence and remote sensing proxies mounted on unmanned aerial vehicles (UAV) or mobile gantries (Herzog et al., 2025). Interest in using spectral datasets has increased in recent years. Spectral information collected from plant tissues such as leaves, stems and clusters, as well as from plant-derived materials including seeds, leaf dry powder or juice can provide a rapid and cost-effective way to characterize plant phenotypes as these spectral profiles are able to capture plant responses to their environment, and serve as complementary omics information layers to support breeding decisions (Robinson et al., 2025a)

Previous studies have primarily focused on the use of leaf optical properties to monitor vine water status (Pampuri et al., 2021), and to model complex physiological traits such as photosynthesis, stomatal conductance, chlorophyll fluorescence or water potential (Rapaport et al., 2015; Coindre et al., 2026). Recently, the use of reflectance signatures from different plant organs has been explored as a low-cost alternative to genomic prediction of agronomic and juice quality-related traits (Brault et al., 2022). Despite these efforts, there is a lack of comprehensive studies that compare how the inherent properties of spectral reflectance from different canopy organs such as leaves and clusters can affect trait prediction across large plant populations.

Hyperspectral information consists of hundreds or thousands of continuous reflectance bands that are high-dimensional and highly collinear. This data structure demands multivariate regression approaches to extract the most information and avoid model overfitting. In this context, the most widely used approach is partial least squares regression (PLSR) as it has shown excellent performance for predicting biochemical, physiological and agronomic traits in annual and perennial crops (Burnett et al., 2021b). Although PLSR is a powerful method for trait prediction, there are still plenty of questions regarding how model performance is associated with the spectral origin of the dataset, the generalization of the models across unknown populations and the minimum number of samples required to build robust models (Burnett et al., 2021a). It remains poorly understood if a given canopy organ will provide with better predictions than other, especially for complex traits that integrate multiple physiological and biochemical processes associated with morphology such as cluster architecture and juice quality.

To address these knowledge gaps, we developed predictive models with reflectance profiles from leaves and clusters to quantify to which extent organ spectra affects model performance and influence the prediction of cluster architecture and juice quality, and two data partitioning strategies were used to understand how the handling of datasets affects trait prediction accuracy and model generalization in two phenotypically diverse Riesling and Pinot clonal populations.

Based on the objectives of this study, we addressed the following research questions: i) Do hyperspectral reflectance profiles differ in their predictive performance depending on the canopy organ from which they derive? ii) Does cluster reflectance provide better prediction accuracy for cluster architecture and juice quality traits compared to leaf reflectance? and iii) How do data partitioning strategies (i.e. cluster type vs population) influence model accuracy and the generalization of trait prediction?

## 2. Materials and Methods

### 2.1 Grapevine populations

This study was conducted during the 2024 harvest season at the experimental vineyards of the Plant Breeding Department at Hochschule Geisenheim University (49°59’11.5“N 7°56’52.4”E). Two extensive clonal populations of *Vitis vinifera* L. were evaluated. These comprised 220 Riesling clones (White Riesling = 217 and Red Riesling = 3) and 240 Pinot clones (Pinot noir = 214, Pinot noir precoce = 7, Pinot blanc = 15 and Pinot gris = 4). Further information regarding each clone, which rootstocks they are grafted on and their field location can be found in (Supplementary Table 1). From this point forward, these populations will be referred as “Riesling” and “Pinot” in the results and discussion sections. These clonal populations have been recently characterised in a multi-year study which demonstrated substantial intra-varietal genetic variability and moderate broad-sense heritability for yield and quality traits in eight commercial grapevine cultivars (Robinson et al., 2025b). Grapevine clusters harvest took place from 3^rd^-9^th^ September for Pinot and 23^rd^-25^th^ September for Riesling, when both populations had reached maturity (BBCH 89) as determined from juice density.

### 2.2 High-throughput phenotyping

From a triplet of the same clone, one healthy cluster was sampled randomly from each vine and grapevine cluster architecture was characterized using a hand-held three-dimensional (3D) scanner (Artec 3D Spider, Senningerberg, Luxembourg) in a lab with controlled light conditions. The images were processed into point clouds using Artec Studio 20 (Senningerberg, Luxembourg) and the 3D bunch tool was used to extract morphological traits including berry number, diameter, volume, and cluster volume, width, length and its convex hull (Rist et al., 2018). Additionally, after scanning, clusters were weighed and spectral signatures were collected.

Cluster reflectance was collected using a portable field spectroradiometer (ASD Field Spec 4, Malvern Panalytical, UK) with a spectral resolution ranging from 350-2500 nm. The reflectance measurements were conducted immediately after the 3D scans under constant light conditions using a leaf clip attachment. The clusters were placed on a black opaque surface to avoid light reflection from the lab bench. Two measurements per cluster were collected on single berries located at opposite sites and averaged to obtain a representative spectral signature for each cluster.

Leaf reflectance was measured with the same device and leaf clip attachment. Three healthy leaves were collected from the sunlit side at the middle of the canopy at flowering (BBCH 65). The leaves were dried in an oven at 60 °C for two days. Then, the leaves were placed on a black opaque surface where spectral signatures were measured under controlled light conditions.

After data collection, raw spectral signatures were filtered using the Savitzky-Golay method to reduce spectral noise (Savitzky and Golay, 1964) due to sensor artifacts or environmental variability using the R package *signal* (Ligges et al., 2015). To smooth the raw signals different window sizes (n) and polynomial orders (p) until visual assessment confirmed the spectral signatures were smoothed, to meet this criterion the values selected in our datasets were n = 11, p = 3. After filtering the signal noise, the spectral range was set to 400-2400 nm as the outer regions of 350-399 nm and 2401-2500 nm were excluded due to low sensor accuracy.

### 2.3 Juice quality analysis

For each of the studied populations, berries from the three replicates per clone were pressed and homogenised to create a composite juice sample. Following pressing, the samples were processed in a centrifuge to clarify the juice and remove supernatants. Then, the juice samples were frozen at -80 °C for later processing. The samples were heated and homogenised in a water bath at 80 °C with regular shaking. The slow warming ensured the dissolution of crystalized sugars and potassium bitartrate (Hieber et al., 2005). Subsequently, the juice was analysed using Fourier-transformed infrared (FTIR) spectroscopy to determine the juice pH, nitrogen compounds assimilable by yeasts (NOPA), juice density and organic acids, including tartaric and malic acid.

### 2.4 Statistical analysis

To build the predictive models we used the best linear unbiased estimators (BLUEs) of reflectance at each wavelength from 400-2400 nm and the 3D phenotypes estimated from the scans. For the quality-determining traits we used the pooled value from the composite sample. BLUEs were calculated per population using the ASReml-R (VSN International Ltd., UK) with the following linear mixed model:

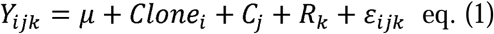

where Y*_ijk_* is the phenotypic observation of Clone*_i_* located at column*_j_* and row*_k_*, μ is the mean, Clone*_i_* is the fixed effect, C*_j_* and R*_k_* are the random effects from the position of the vine in the field and ε*_ijk_* is the residual error. Random effects (C*_j,_* R*_k_*) and residuals were assumed to be independently and identically distributed with mean zero and constant variance.

Trait differences among populations were evaluated using principal component analysis (PCA) to visualize the extent of separation between populations. To determine which traits differed significantly between the Riesling and Pinot populations, a multivariate analysis of variance (MANOVA) with population included as fixed effect was used.

### 2.4.1 Partial least squares regression modelling

Partial least squares regression (PLSR) was used to model the associations between hyperspectral signatures (X) with cluster architecture and quality traits (Y). This supervised multivariate regression approach identifies latent variables that maximize shared covariance between highly dimensional predictors and the response variable (Boulesteix and Strimmer, 2006). PLSR decomposes X and Y as follows:

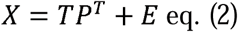

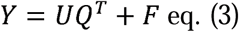

where X is the predictor variable, Y is the response, T and U are the latent scores summarizing the covariance structure of X and Y, respectively; P and Q are the loading coefficients for the contribution of each predictor to the response variable, E and F are the residual matrices.

Furthermore, the latent variables are obtained maximizing the covariance between X and Y projections as follows:

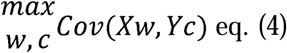

where w and c are the weight vectors defining the latent directions of X and Y, respectively. Finally, the regression models can be expressed as:

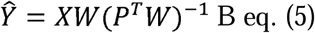

where W is the weight matrix and B is the regression coefficient of the latent structure relating X to Y.

Variable importance in projection (VIP) scores were calculated from the calibrated PLSR models to identify the spectral regions contributing the most to trait prediction by integrating the PLS variable weights across latent components and the response variance explained from each component.

PLSR trait predictions were implemented with the orthogonal scores PLS algorithm from the *pls* package in R (Mevik and Wehrens, 2007). Hyperspectral predictors consisted of matrices from i) dry leaf reflectance, ii) cluster reflectance and we included four synthetic spectral signatures created with different matrix operations, these were iii) averaged organ reflectance, iv) combined organ reflectance, v) reflectance matrix prioritising cluster spectra importance (0.7 x cluster reflectance value, 0.3 x dry leaf reflectance value: 70% cluster/30% dry leaf; from now on 70_30) and vi) reflectance matrix prioritising dry leaf spectra importance (0.3 x cluster reflectance value, 0.7 x dry leaf reflectance value: 30% cluster/70% dry leaf; from now on 30_70).

To evaluate how data partitioning strategies influence model predictions, two independent approaches were implemented. In the first approach, observations were grouped according to cluster architecture, following the International Organisation of Vine and Wine classification for bunch density (OIV descriptor 204), which categorizes the clusters as compact, intermediate or loose. Based on this classification, the Riesling population comprised 74 compact, 109 intermediate and 37 loose cluster phenotypes; whereas the Pinot population included 68 compact, 131 intermediate and 41 loose cluster phenotypes (Supplementary Table 1).

For the second approach, observations were grouped according to population origin, distinguishing between the Riesling and Pinot clones. This, predictive models were developed under two data partitioning schemes: i) cluster architecture-based models and ii) population-based models.

Within each partitioning strategy, the datasets were randomly split into a training set (80%) used for model calibration and an independent validation set (20%) used for model evaluation. PLSR models were calibrated using the training dataset to find the optimal number of latent components using a cross-validation (CV) approach within the training set by selecting the number of components that minimised both the root mean square error (RMSE) and the predicted residual sum of squares as previous studies have suggested (Serbin et al., 2014; Robles-Zazueta et al., 2022). Finally, model performance was assessed on the independent validation set using the determination coefficient (R^2^), RMSE and model bias (Supplementary Table 3).

## 3. Results

### 3.1 Phenotypic variation across populations and cluster types

Our results revealed variation among cluster architecture types and between the Riesling and Pinot populations. Across both populations, compact clusters exhibited a higher berry number, greater cluster volume, and higher cluster weight compared with intermediate and loose clusters (Figure 1A, 1C, 1D, 1E, 1G, 1H). In addition, a general trend was observed in which larger clusters were associated with both a higher berry number and larger berries (Supplementary Figure 1).

**Figure 1.**
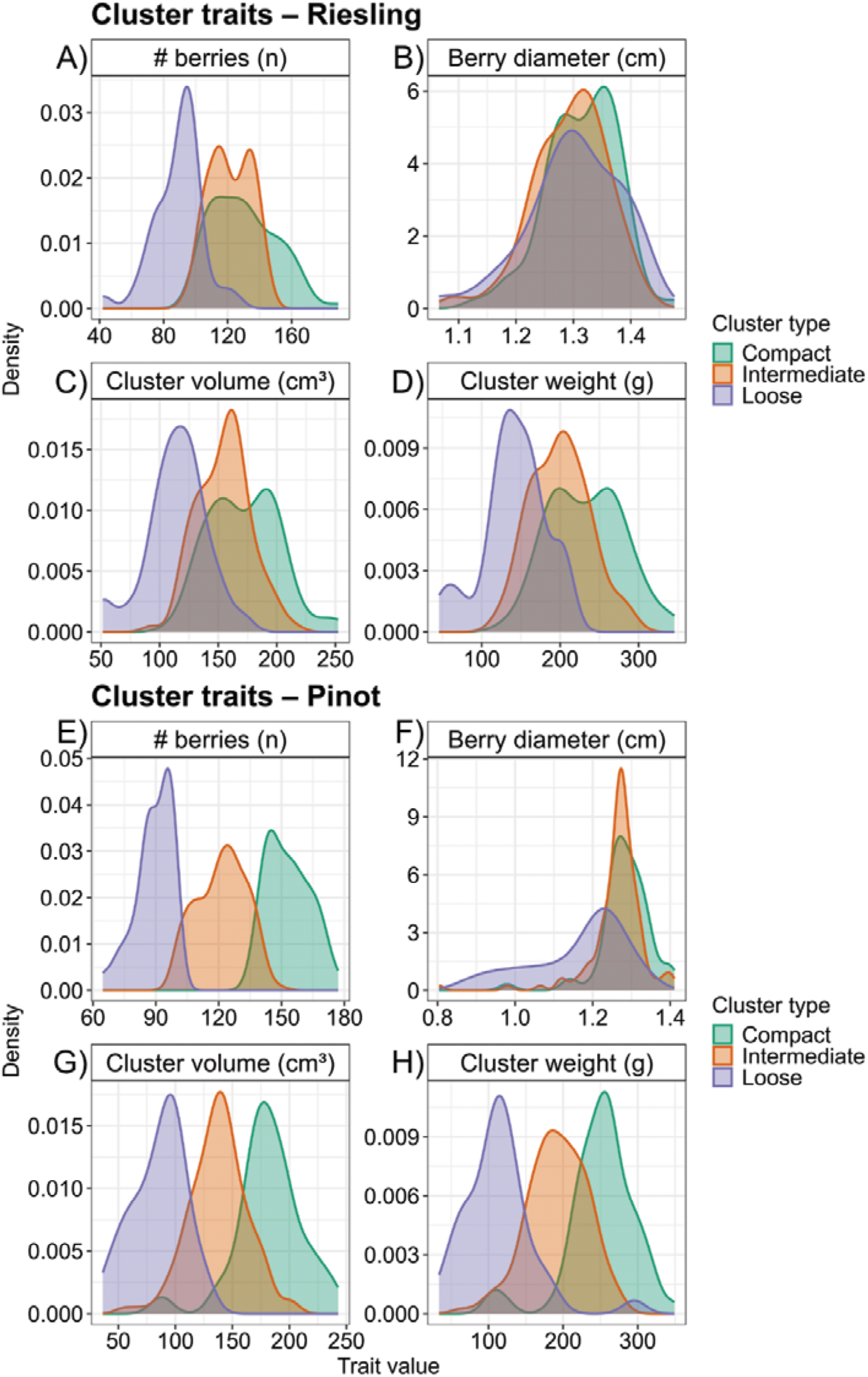
Density plots of cluster traits distribution in Riesling and Pinot clonal populations. The colours from the curves represent three distinctive cluster architecture types based on the OIV classification (compact = green, intermediate = orange, loose = purple). Panels A-D) show the distribution of cluster traits in the Riesling population and panels E-H) show the distribution of cluster traits in the Pinot population.

Principal component analysis further indicated a clear separation between the Riesling and Pinot populations, whereas the three cluster architecture types largely overlapped within each population (Supplementary Figure 2). Consistent with this pattern, Riesling clusters were significantly larger in volume (Riesling: 154.60 ± 32.06 cm^3^, Pinot: 138.30 ± 41.05 cm^3^; p<0.001), contained more berries (Riesling: 176 ± 30, Pinot: 123 ± 24; p<0.001) and had larger berry size (diameter = 1.57 ± 0.04 cm, Pinot: 1.25 ± 0.09 cm; p<0.001, Figure 1B) compared to Pinot clusters (Figure 1F). In contrast, cluster architecture traits, including convex hull, cluster length and width did not differ significantly between populations (Supplementary Table 2).

For juice quality traits, Riesling showed significantly lower pH values (2.91 ± 0.08) compared to Pinot (3.15 ± 0.14; p<0.001, Supplementary Table 2). Consistent with this pattern, Riesling also exhibited higher total acidity (Figure 2A, Figure 2D) and tartaric acid concentrations compared to Pinot (p<0.001; Supplementary Table 2). In contrast, Pinot showed higher concentrations of NOPA (141.17 ± 47.37 mg N/L, p<0.001; Figure 2F) and total sugars (184.09 ± 14.22 g/L, p<0.001; Figure 2E, Supplementary Table 2) compared to Riesling (NOPA: 75.28 ± 47.37 mg N/L; total sugars: 172.20 ± 15.31 g/L).

**Figure 2.**
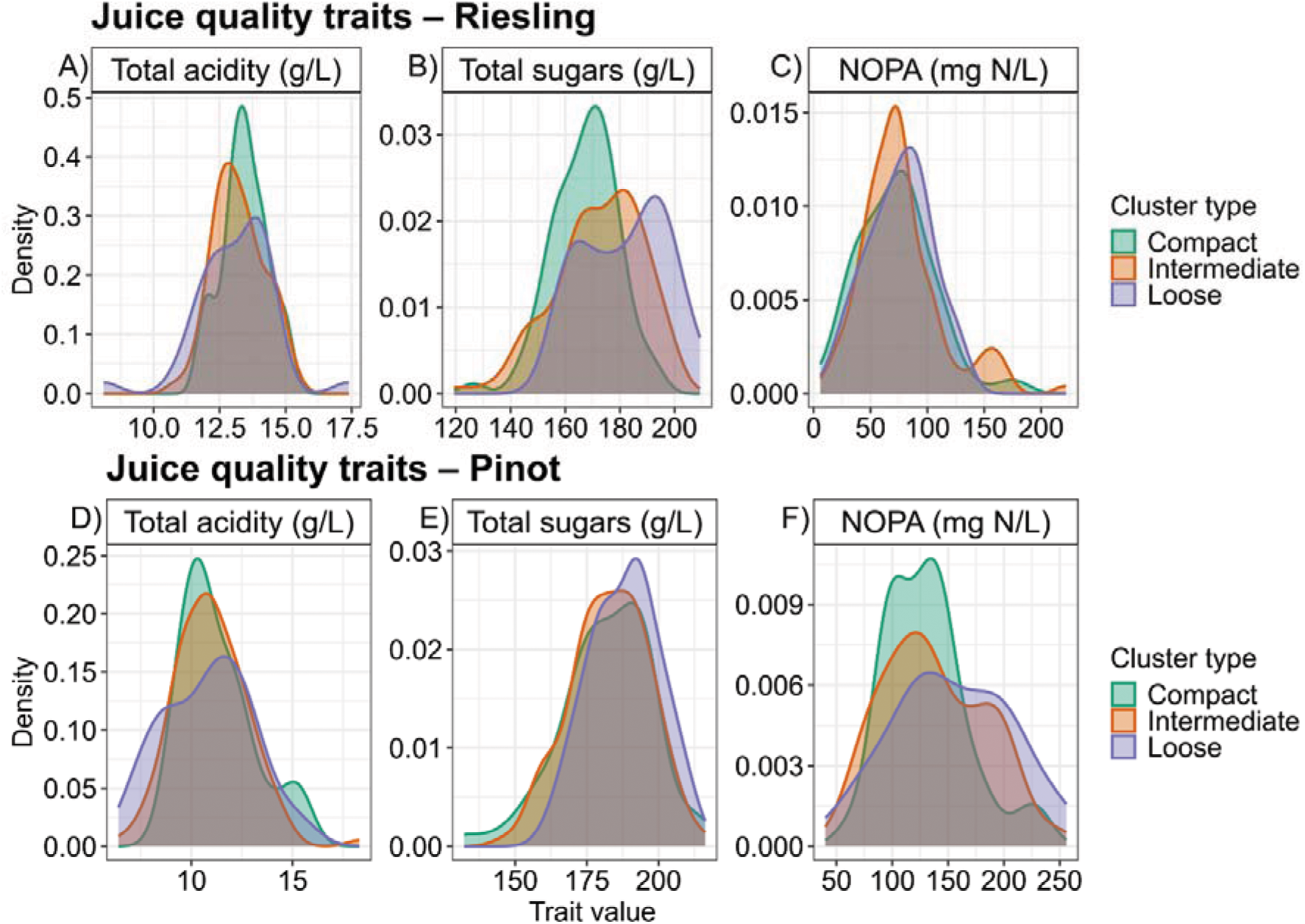
Density plots of juice quality traits distribution in Riesling and Pinot clonal populations. The colours from the curves represent three distinctive cluster architecture types based on the OIV classification (compact = green, intermediate = orange, loose = purple). Panels A-C) show the distribution of juice quality traits in the Riesling population and panels D-F) show the distribution of juice quality traits in the Pinot population.

Within the Riesling population, malic (2.8-9.4 g/L, Supplementary Table 2) and tartaric acid (4.4-10 g/L, Supplementary Table 2) concentrations showed relatively similar distributions across cluster architecture types, whereas total acidity (Figure 2A), total sugars (Figure 2B), and NOPA (Figure 2C) exhibited larger variability, particularly for intermediate and loose phenotypes. In contrast, the Pinot population showed greater variation in malic, tartaric acid concentrations and NOPA (Figure 2F), while pH and total sugars (Figure 2E) were more uniformly distributed across cluster architecture types. Overall, the trait phenotypic variation showed the higher acidity typically associated with Riesling juice (Supplementary Table 2).

Hyperspectral reflectance profiles from dry leaves showed distinct spectral patterns between populations (Figure 3A, 3B) In both populations, reflectance spectra were low in the visible (VIS: 400-700 nm) region, consistent with strong light absorption from chlorophyll, carotenoids and anthocyanins. In the Pinot population, a sharp increase in reflectance was observed in the red-edge region (∼680-720 nm, Figure 3B). In contrast, Riesling showed more variability across red-edge and near-infrared (NIR) regions, as well as across the shortwave infrared (SWIR) area in comparison to Pinot (Figure 3A). Both populations exhibited pronounced absorption features associated with water molecules at ∼1200 nm and between 1400-1900 nm.

**Figure 3.**
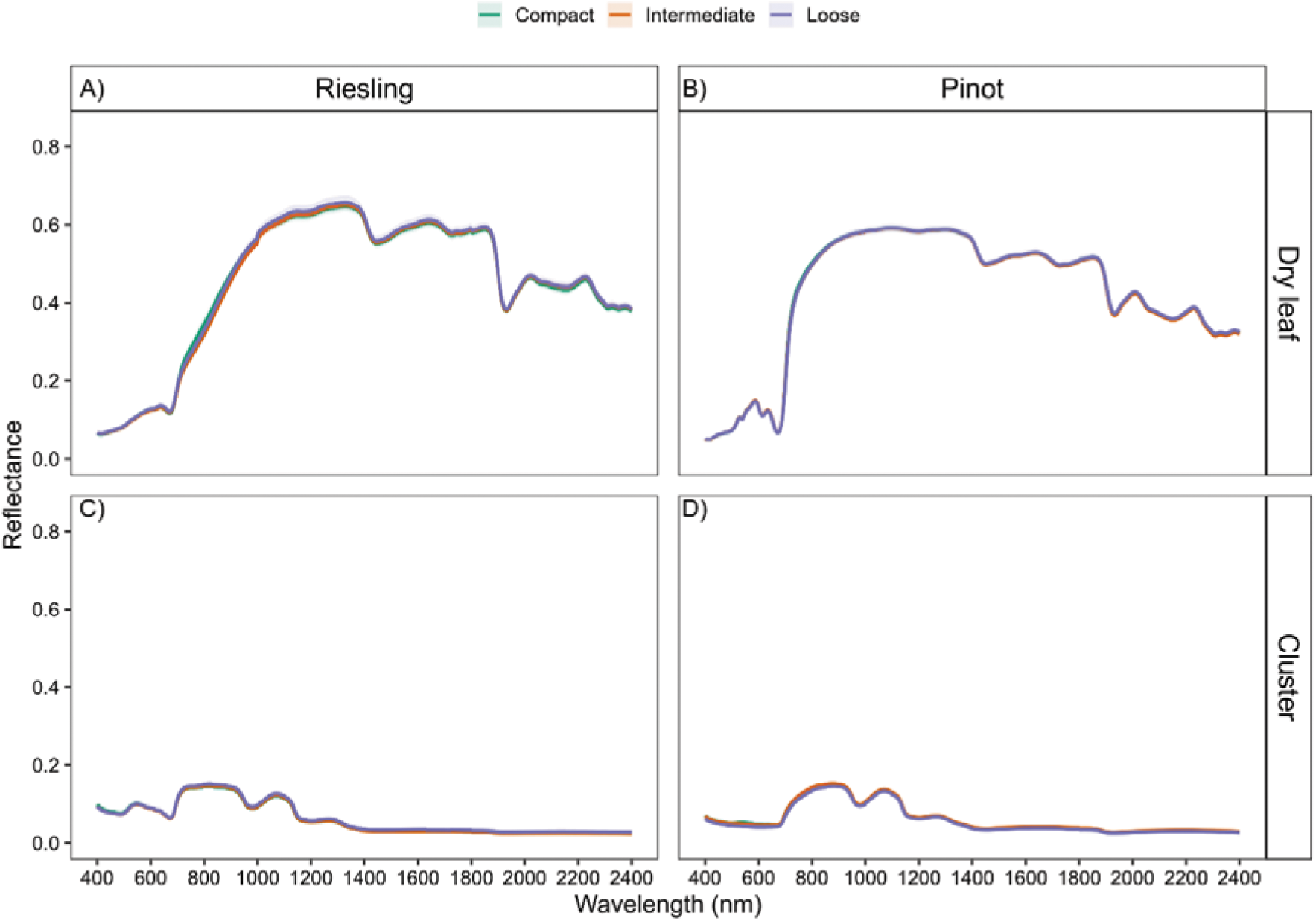
Mean hyperspectral reflectance profile of dry leaves and clusters of Riesling and Pinot clones. Reflectance profiles of dry leaves (A, B) and grapevine clusters (C, D) are shown for each population. Coloured lines indicate the cluster architecture types (compact = green, intermediate = orange, loose = purple).

In contrast, cluster reflectance profiles were more uniform between populations (Figure 3C, 3D). The high-water content present in berries resulted in very low reflectance values (<0.1) across the SWIR region. In addition, the absence of photosynthetic pigments in berries led to lower reflectance in the VIS and NIR regions compared with leaf spectral profiles.

These population trait differences provided the basis for evaluating predictive models testing different data partitioning strategies and spectral profiles.

### 3.2 Predictive modelling within population and cluster architecture groups

Model performance for cluster architecture traits varied depending on the trait predicted, the reflectance profile used for model calibration, and the data partitioning strategy applied. When cluster reflectance was used as predictor, models estimating berry traits achieved higher predictive accuracy than those predicting cluster traits. For example, berry diameter was predicted with relatively high accuracy (R^2^ = 0.76, RMSE = 0.09 cm), whereas predictions for cluster weight (R^2^ = 0.06, RMSE = 61.48 g), cluster volume (R^2^ = 0.05, RMSE = 37.97 cm^3^) and cluster convex hull (R^2^ = 0.09, RMSE = 67.18 cm^3^) showed considerably lower accuracy (Figure 4). In general, models trained on datasets partitioned by cluster architecture types showed higher validation R^2^ and, in several cases, lower RMSE than models split by population, although this pattern was trait- and predictor-dependent (Supplementary Table 3).

**Figure 4.**
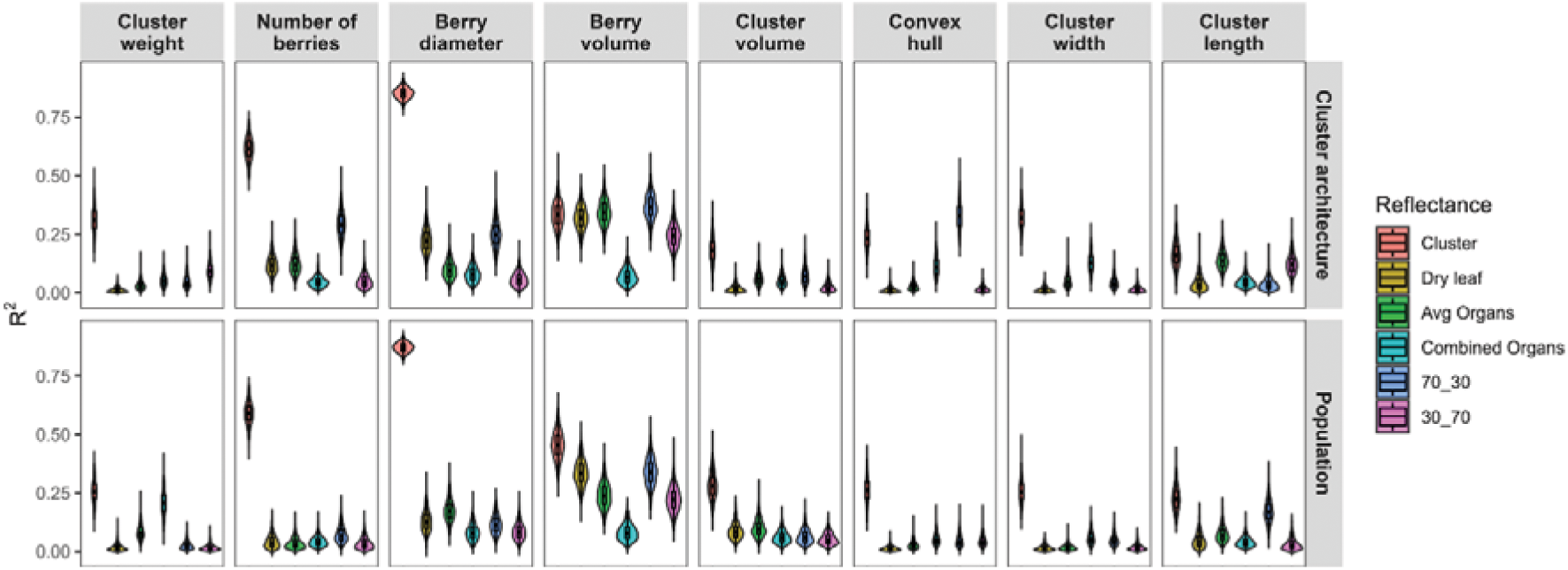
Violin plots of the model training for cluster architecture traits. The plot shows the R^2^ from the model trained with different reflectance profiles such as cluster spectra (orange), dry leaf spectra (brown), averaged organs (green), combined organs (cyan), 70_30 (navy blue) and 30_70 (pink). The panels are divided by dataset partitioning strategies.

The spectral profiles used for model calibration showed a trait-dependent effect on prediction accuracy. For cluster architecture traits, cluster reflectance-based models frequently produced lowest RMSE, although differences among spectral profiles were small (Supplemental Figure 3). Dry leaf and synthetic reflectance profiles had comparable performances across cluster architecture and juice quality traits, whereas models based on combined spectral profiles tended to produce the less accurate models (Supplementary Figure 3, 4).

Among cluster architecture traits, the highest prediction accuracies using cluster reflectance were obtained for berry number (calibration R² = 0.52, RMSE = 26 berries per cluster; CV R² = 0.44, CV RMSE = 28 berries per cluster) and berry diameter (calibration R² = 0.82, RMSE = 0.07 cm; CV R² = 0.79, CV RMSE = 0.08 cm). Predictions for other cluster traits were generally lower, although cluster reflectance models still tended to produce higher R² and lower RMSE than the other spectral profiles (Supplementary Table 3).

On the other hand, predictive models for juice quality traits showed higher accuracy than those for cluster or berry traits. Total acidity (R² = 0.48, RMSE = 1.42 g/L) and pH (R² = 0.63, RMSE = 0.10) were predicted with relatively high accuracy, particularly when models were trained using cluster-type partition and calibrated with dry leaf reflectance (Figure 5). Overall differences in prediction accuracy from data partitioning strategies were relatively small. Model performance varied more strongly with the spectral profile used for calibration, with several cluster architecture traits (Figure 4) and a few juice quality traits (Figure 5) showing clear differences in R^2^.

**Figure 5.**
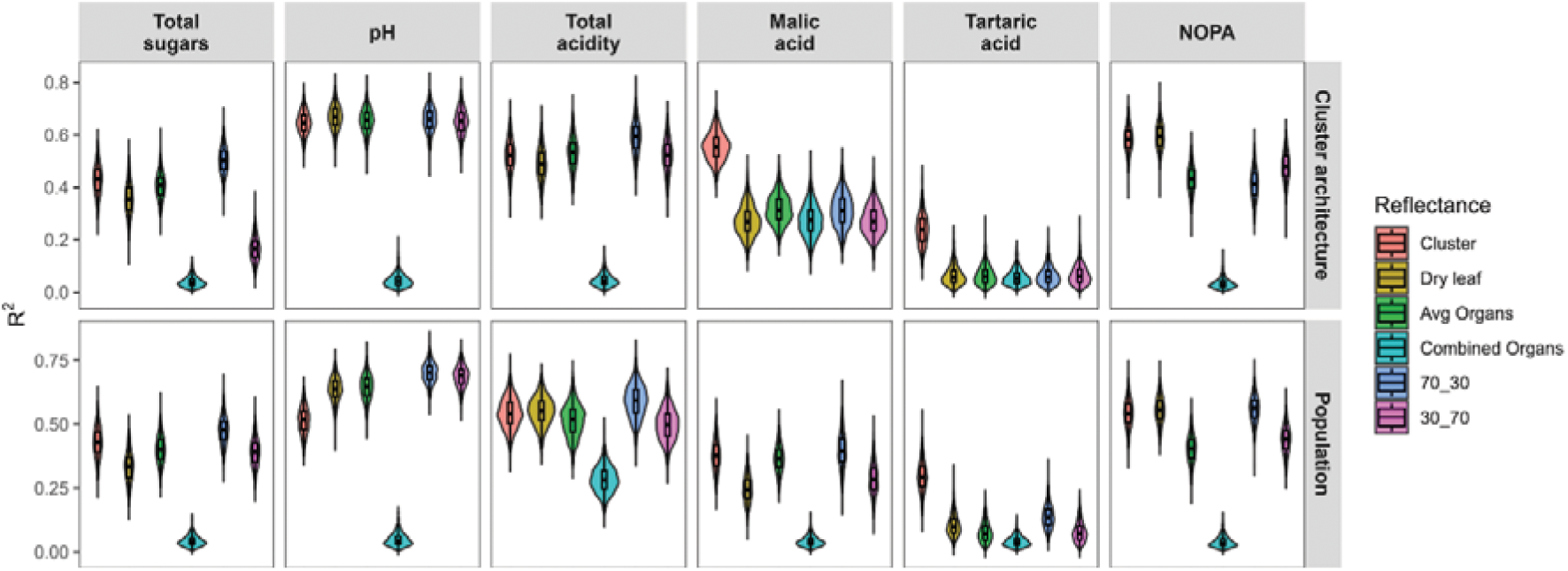
Violin plots of the model training for juice quality traits. The plot shows the R^2^ from the model trained with different reflectance profiles such as cluster spectra (orange), dry leaf spectra (brown), averaged organs (green), combined organs (cyan), 70_30 (navy blue) and 30_70 (pink). The panels are divided by dataset partitioning strategies.

Given the relatively small effects found due to data partitioning strategies, the next step was to examine how the spectral profiles used for model calibration can influence prediction accuracy.

### 3.3 Influence of spectral profiles on trait prediction accuracy

To further examine how spectral profiles affected model performance, we evaluated their predictive accuracy across architecture and juice quality traits in the validation dataset. Results showed that the prediction accuracy of PLSR models depended strongly on the spectral profile used for calibration and the trait predicted. Overall, cluster architecture traits (Figure 6) were more difficult to predict than juice quality traits (Figure 7).

**Figure 6.**
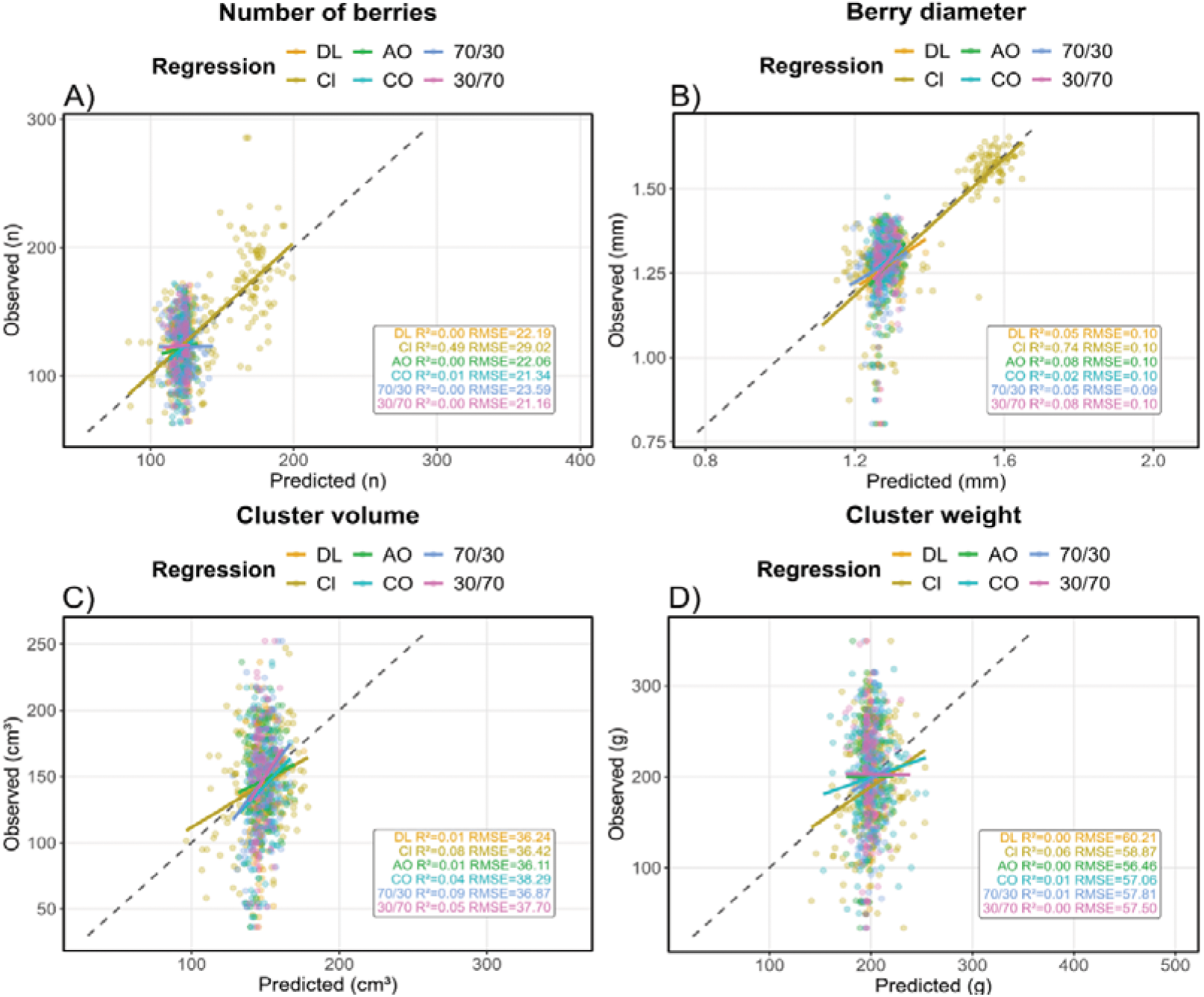
PLSR model validation regression plots for cluster architecture traits. Different colours indicate the spectral profile used as input to validate the model. DL = dry leaf, Cl = Cluster, AO = Averaged organs, CO = Combined organs, 70/30: 70% importance dry leaf, 30% importance cluster, 30/70: 30% importance dry leaf, 70% importance cluster. A) number of berries, B) berry diameter, C) cluster volume and D) cluster weight. The solid lines represent the fitted regression for each spectral input and dashed lines show the 1:1 relationship.

**Figure 7.**
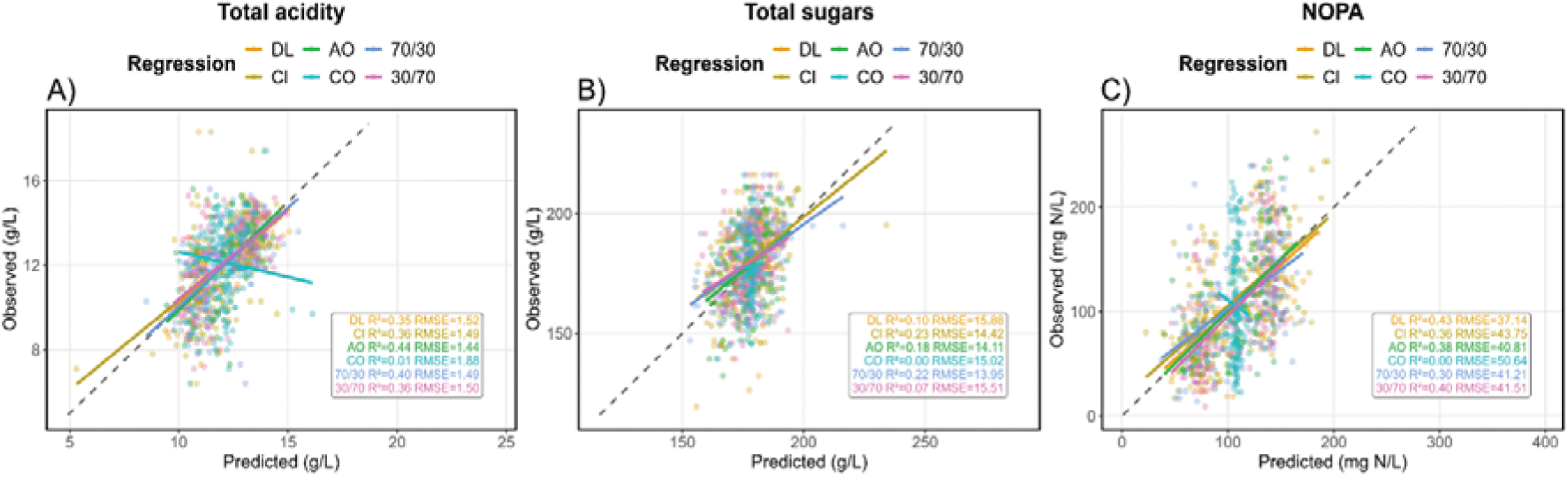
PLSR model validation regression plots for quality traits where each point represents an individual clone measurement and the different colours indicate the spectral profile used as input to validate the model. DL = dry leaf, Cl = Cluster, AO = Averaged organs, CO = Combined organs, 70/30: 70% importance dry leaf, 30% importance cluster, 30/70: 30% importance dry leaf, 70% importance cluster. A) total acidity, B) total sugars and C) NOPA. The solid lines represent the fitted regression for each spectral input and dashed lines show the 1:1 relationship.

Berry number and diameter were predicted with high accuracy when cluster spectra was used, outperforming models calibrated with dry leaf or synthetic spectral profiles (Figure 6A, 6B). Berry volume showed a similar pattern but with lower overall accuracy (Supplementary Table 3). Predictions for cluster-level traits such as weight and volume were generally poor across all spectral profiles (Figure 6C, 6D).

In the juice quality traits, the predictive accuracy was generally higher compared to the cluster architecture traits and varied on the different spectral sources. The highest accuracy found in this study was obtained for pH predictions using dry leaf reflectance (R^2^ = 0.62, RMSE = 0.1; Supplementary Table 3). For total acidity and NOPA, predictive performance was comparable across all spectral profiles, except for the average organ reflectance, while for total sugars the models built with cluster reflectance had the best performance. Despite these differences among spectral profiles, some quality traits remained difficult to predict, with malic and tartaric acids showing low predictive accuracy across reflectance sources (Supplementary Table 3).

Overall, these results highlight the selection of spectral profiles depend on the trait of interest. Cluster reflectance profiles are more informative to predict berry traits, whereas dry leaf and synthetic spectral profiles are comparable or have better performance for juice quality traits.

### 3.4 Key spectral regions contributing to trait prediction

Variable importance in projection (VIP) scores were used to identify the spectral regions contributing most strongly to the PLSR models. When models were calibrated using dry leaf reflectance, the most important wavelengths were found in the VIS, red-edge and NIR regions of the spectrum. For cluster architecture traits, the peaks were found in the red-edge and NIR regions (Figure 8A). Specifically, the spectral regions 718-868 nm, 698-868 nm and 654-978 nm contributed the most to the predictions of berry number, diameter and volume, respectively (Table 1).

**Figure 8.**
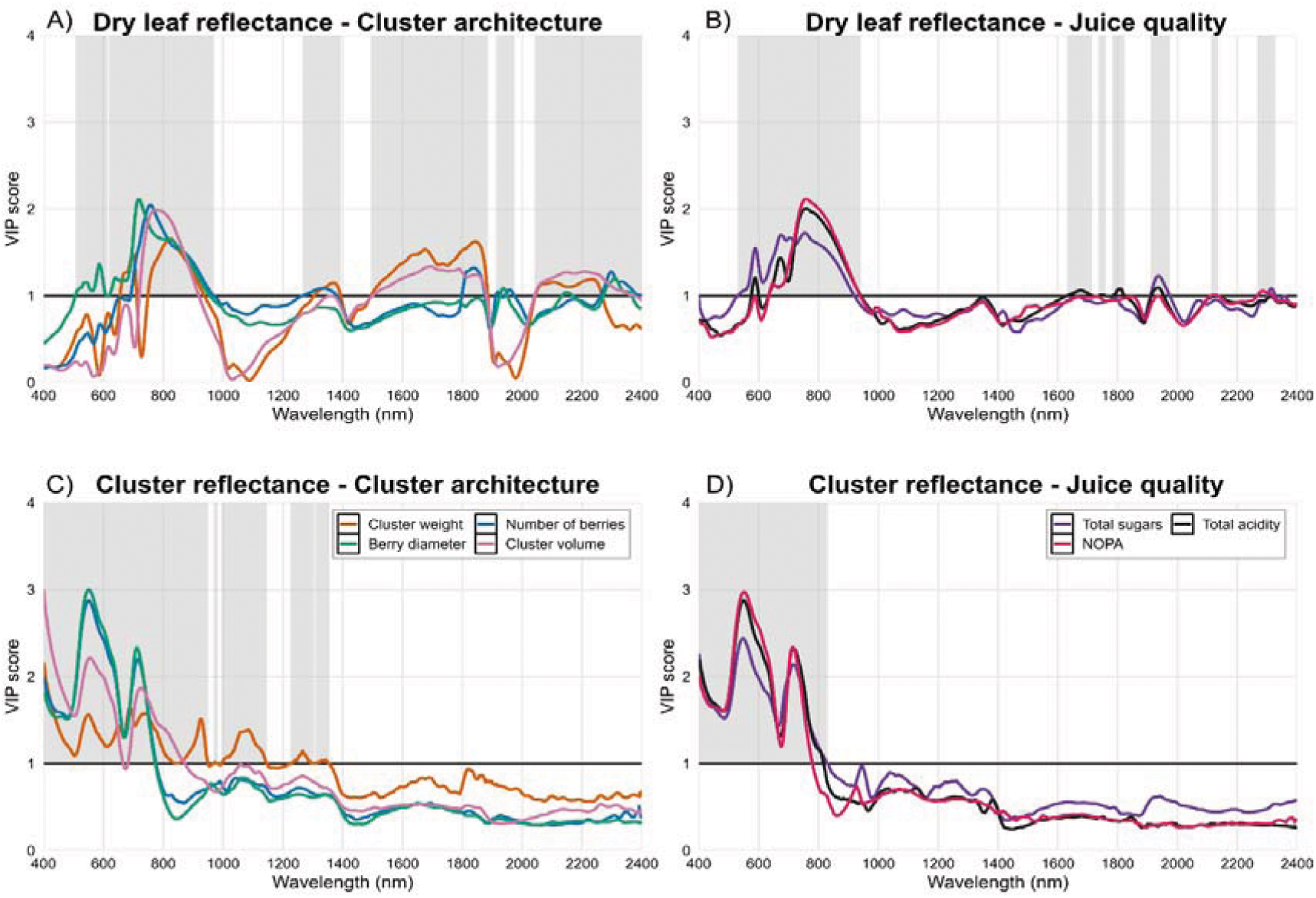
Variable importance in projection (VIP) scores to predict cluster architecture (A, C) and quality traits (B, D) across populations. VIP scores across the visible (400-700 nm), NIR (700-1300 nm) and SWIR (1300-2400 nm). Panels A and B are scores derived from the use of dry leaf spectra as predictor, while panels C and D use cluster spectra. The solid black line is the VIP threshold above which wavelength reflectance is considered to be important for trait prediction, and the grey bands represent areas of the spectra with importance for the prediction of the traits.

**Table 1.** Spectral wavelengths with greater importance for trait prediction for the two clonal grapevine populations based on variance importance in projection (VIP) scores >1.

| Trait | Dry leaf reflectance | Cluster reflectance |
| --- | --- | --- |
| Cluster weight (g) | 695-705; 790-856; 1662-1691; 1792-1868 | 400-468; 524-830; 810-939; 1048-1143; 1211; 1216; 1221-1227; 1233-1276 |
| Berry number (#) | 718-868 | 400-775 |
| Berry diameter (cm) | 587-588; 698-868 | 400-772 |
| Berry volume (cm<sup>3</sup>) | 584-592; 654-978; 991-993; 996; 1691-1740; 1908-1968 | 400-764 |
| Cluster volume (cm<sup>3</sup>) | 726-889; 1666-1740; 1787-1798 | 400-867; 870-873; 1048-1051; 1053; 1057; 1091; 1093-1094 |
| Cluster convex hull (cm<sup>3</sup>) | 702-859 | 400-487; 512; 522-809; 906; 918; 921; 923; 925; 927-932; 939-943; 950-995; 1028-1057; 1066; 1097; 1103-1122; 1139; 1159; 1328; 1804-1828; 1898; 1910-1961 |
| Cluster width (cm) | 526-907 | 400-437; 447; 508; 511-808; 814; 820; 827; 830; 906; 911; 916; 920-999; 1004-1130; 1138-1163; 1177; 1186; 1241; 1328-1336; 1368; 1386; 1412-1413; 1424; 1492-1499; 1519; 1813-1815; 1897-1983; 2381-2397 |
| Cluster length (cm) | 712-882; 1615-1716 | 400-657; 696-945; 1010-1133 |
| Total sugars (g/L) | 528-919; 1792-1797; 1911-1974; 2303-2362 | 400-828; 831-833; 937-939; 944-945; 951; 1042; 1044 |
| pH | 582-595; 638-944; 1685-1692; 1701-1705; 1796-1797; 1908-1965; 2114-2139; 2312-2313 | 400-784 |
| Total acidity (g/L) | 577-598; 634-940; 1632-1712; 1737-1761; 1779-1782; 1790-1798; 1802-1819; 1910-1953; 2308-2316 | 400-817 |
| Malic acid (g/L) | 576-603; 630-652; 655-748; 1804-1805; 1925-1951 | 400-998; 1023-1105; 1107-1108; 110-1128; 1140-1159; 1210-1256; 1261; 1270; 1275-1278; 1281; 1284; 1291 |
| Tartaric acid (g/L) | 712-918 | 400-841; 845-846; 1002-1022; 1140-1141; 1145; 1370; 1376; 1380 |
| NOPA (mg N/L) | 688-913 | 400-778 |

A similar spectral pattern was observed in the juice quality traits when using dry leaf reflectance (Figure 8B). The highest VIP scores were located in the VIS and red-edge regions, particularly for total sugars, total acidity and malic acid, which showed large peaks between 638-944 nm, 634-940 nm and 630-748 nm, respectively (Table 1). In contrast, pH, tartaric acid and NOPA showed stronger contributions from wavelengths located in NIR water absorption regions (Table 1).

Models calibrated using cluster reflectance profiles, exhibited that the most important wavelengths to predict cluster architecture are located between 400-1100 nm, with number of berries, berry diameter and cluster volume showing VIP scores >2 (Figure 8C). Additional peaks observed in the NIR and SWIR regions between 950-1200 nm contributed to the prediction of cluster-related traits (Table 1). The juice quality traits, concentrated the most important spectral regions in the VIS and red-edge regions (Figure 8D). With the most influential wavelengths located between 400-778 nm (Table 1).

Overall, prediction accuracy varied across traits, with berry and juice quality traits generally showing higher accuracy than cluster-level architecture traits. Model performance was more strongly associated with the spectral profile used for calibration than with the data partitioning strategy. VIP results indicated that wavelengths contributing most to trait prediction were concentrated in the VIS and red-edge with additional contributions from NIR water absorption bands.

## 4. Discussion

Hyperspectral reflectance provides a framework to develop robust phenomic proxies for complex agronomic and quality traits in breeding programs. However, the prediction accuracy of the models depends on how datasets are structured and the biological source of the reflectance profile. In this study, we evaluated how data partitioning strategies and reflectance profiles from different plant organs influence the prediction of cluster architecture and juice quality traits in two populations of Riesling and Pinot clones. Our findings emphasize that prediction accuracy is trait-dependent and that the source of spectral profiles substantially influence phenomic models performance.

### 4.1 Population variation for cluster architecture and juice quality traits

The phenotypic variation between Riesling and Pinot clonal populations provide important context for interpreting the predictive models in our study. Compact cluster types in both populations frequently combined high number of berries with larger cluster volume and weight. These cluster patterns are consistent with previous evidence showing that berry number, berry diameter, cluster weight and cluster length can explain >50% of bunch architecture variability (Rist et al., 2022). However, there are certain cultivar clones exhibiting small and low weight clusters that have numerous small berries that can be compact due to the short pedicel lengths in their rachis (Rossmann et al., 2020). These examples show the limitations of non-invasive remote sensing approaches where traits which are subject to occlusion may affect model prediction accuracy.

In our populations, Pinot showed stronger modulation of berry diameter and volume across cluster architecture types, supporting previous findings that cluster architecture is a set of traits strongly dependent on the genetic background of the population (de Oliveira et al., 2025; Richter et al., 2019). From a viticulture perspective, loose bunch architecture is a major breeding target because it reduces the risk of Botrytis bunch rot, minimizes berry compression that promotes splitting during ripening and overall improves the bunch microclimate.

Cluster architecture is indirectly associated with juice quality mainly through its effects on berry size. Higher total acidity and lower pH were found to be linked to loose cluster types due to improved bunch aeration and reduced malate respiration, whereas compact types often were less acidic. However, soluble sugar content in the bunches was not consistently associated with bunch architecture (Frioni et al., 2023; Nistor et al., 2025). Evidence linking cluster architecture with NOPA remains limited, as the environmental conditions and nutritional status of the vine play a stronger role than bunch architecture (Marcuzzo et al., 2025). Together, these findings support the broader concept that cluster architecture is indirectly controlling juice quality through microclimatic regulation in the bunch (Keller, 2020).

The contrasting distribution of organic acids across cluster types observed among Riesling clones, compared with the more uniform ranges of the Pinot clones, further highlights the population-specific nature in these relationships. This observation is consistent with previous multiyear analysis in these clonal populations (Robinson et al., 2025b) and with genomic studies that have demonstrated that berry quality traits and their predictability are strongly genotype dependent (Flutre et al., 2023).

### 4.2 Effects of data partitioning strategies on trait predictability

Our modelling results showed that the data partitioning strategy had a low influence on prediction accuracy in comparison with the spectral profile used for model calibration. Models trained using cluster-type partitions capture more variability in cluster traits but generally caused slightly higher accuracy variability in the model validation. In contrast, the population-based partitioning resulted in lower predictive accuracy likely because grouping observations by population reduced the phenotypic and spectral variability available for model calibrations.

This situation resembles many phenomic studies in grapevine where models are developed using data from a single cultivar, environment or management conditions. Similar approaches used to predict vine physiological traits have used leaf reflectance from the SWIR to predict stem water potential and leaf relative water content with high correlations ranging from 0.66-0.81 (Tardaguila et al., 2017). Pre-dawn water potential has also been predicted in a small Pinot blanc population (n = 24) with leaf spectral profiles from 350-2500 nm achieving a prediction accuracy of R^2^ = 0.7 (Pampuri et al., 2021), and transpiration (R^2^ = 0.66) and stomatal conductance (R^2^ = 0.66) was predicted in a small population (n = 36) of Syrah, Riesling and Merlot cultivars (Ryckewaert et al., 2022).

Our observation that dataset partitioning can influence prediction accuracy and model stability is consistent with previous studies, although depending on the context. For example, predictions of stem water potential were lower when the models used a simple cultivar (Tempranillo, R^2^ = 0.79) compared to when the models combined multiple cultivars (R^2^ = 0.84, Gutierrez et al., 2016). Similarly, data partitioning strategies affected the prediction of leaf physiological and biochemical traits across a set of diverse cultivars including Cabernet Franc, Cabernet Sauvignon, Merlot, Pinot noir, Sauvignon blanc and Viognier, where random data partitioning improved prediction of photosynthetic traits, while cultivar-based partitioning improved prediction of biochemical traits (Cui et al., 2025). Comparable patterns have been reported in other crops, where phenomic models trained on genetically and environmentally diverse datasets captured more trait variability and improved trait prediction for biomass, leaf water content, photosynthesis and chlorophyll fluorescence (El-Hendawy et al., 2020; Robles-Zazueta et al., 2022).

Phenomic models developed on narrow or homogeneous germplasm (i.e. single cultivars or environments) often show limited transferability because they capture a restricted range of spectra-trait covariance (Grzybowski et al., 2021). In contrast, models calibrated with diverse germplasm and environments tend to be more generalizable, improving the predictive ability and heritability of phenomic trait predictions (Galán et al., 2021). Together, these findings reinforce a core principle for genomic and phenomic prediction approaches that calibration datasets which capture broad phenotypic and genetic diversity are essential for building transferable predictive models.

### 4.3 Importance of different organ reflectance and spectral regions

One of the main findings in this study was that prediction accuracy depends on the spectral profile used for model calibration. The VIP scores revealed that spectral regions and plant organ from which reflectance was measured strongly determines model performance.

Cluster reflectance provided the most informative signals for several cluster and some juice quality traits, with peaks located in the VIS and red-edge regions. These wavelengths are related to anthocyanin, carotenoid and flavonoid light absorption present in the berry skins. Previous studies have shown that spectral indices derived from VIS and NIR regions are able to capture variation in berry anthocyanin content (Blank and Stoll, 2019), as well as associations with sugar accumulation at ripening in Riesling and Cabernet Sauvignon (Navrátil and Buschmann, 2016). Further, hyperspectral reflectance models have achieved high prediction accuracies for glucose (R^2^ = 0.71), fructose (R^2^ = 0.64) and malate (R^2^ = 0.55) of individual berries across the ripening period (Tavernier et al., 2025). In contrast, dry leaf reflectance showed lower predictive ability for most cluster architecture traits. Dry leaf spectra primarily reflect structural carbon compounds, residual pigment content and internal light scattering structures (Jacquemoud and Ustin, 2019), which is likely to not be related to cluster morphology or berry composition.

The higher prediction accuracy of pH from dry leaf spectra may reflect physiological coupling between leaves and berries through ion regulation. Potassium and calcium content increase during ripening and contribute to the osmotic adjustment and acid neutralization in berries (Rogiers et al., 2017), potentially linking leaf spectral features to juice pH.

Dry leaf and dry wood reflectance profiles have been previously used to predict agronomic and quality traits with greater accuracy compared to when the profiles were used individually in a diverse grapevine population consisting of commercial varieties and breeding lines (Brault et al., 2022). However, our results indicate that mixing spectral profiles from different organs may reduce prediction accuracy, especially as the organs differ in biochemical and structural compositions which can dilute trait-specific associations, or if data collection occurs at different developmental stages. Similar organ-specific effects have been reported where leaf reflectance models predicted berry quality with higher accuracy than berry reflectance models across 23 grapevine genotypes (Wong et al., 2025), and radiative transfer models have demonstrated that VIP signals vary depending if the predicted traits are related to plant nutrition, biochemistry or canopy structure (Farajpoor et al., 2025).

Despite the promising prediction results obtained in this study, hyperspectral phenomic models present inherent limitations related to the spatiotemporal variability in reflectance. Spectral signatures vary with phenological stages, canopy structure and environmental conditions, which affects model generalization across seasons or environments. Additionally, grapevine trials in the field (and most perennial tree crops) often involve logistical constraints that lead to unbalanced experimental designs with limited replication. In our study, the clonal populations were distributed across different vineyard blocks and the dataset was not fully balanced across populations and cluster architecture types, meaning that genetic variability could be partially confounded with environmental effects. Consequently, model performance should be interpreted with caution and future studies should aim to evaluate phenomic models across replicated vineyard sites with multiple seasons, broader genetic and phenological variability.

## Conclusions

This study demonstrates that organ-specific hyperspectral reflectance can provide meaningful phenomic predictions for cluster architecture and juice quality traits in grapevine. Prediction accuracy varied among traits, with berry and juice quality traits showing higher predictive accuracy compared to cluster-level traits. Our results indicate that the spectral profile used for model calibration is more important than the data partitioning strategy, highlighting the importance of selecting the appropriate plant organ when designing phenomic models. In particular, cluster reflectance captured variation in cluster architecture traits more effectively, and our analysis revealed that the predictive information was concentrated in the VIS, red-edge and NIR water absorption bands.

Future studies in perennial crops should focus on improving model foundation to integrate spectral information from multiple plant organs and phenological stages. Combining reflectance profiles from sunlit and shaded leaves, clusters, stems, as well as canopy-level observations could better capture the physiological and biochemical features present in spectral information to underline the spectra-trait associations in grapevine. The key spectral regions found in our study provide a foundation for spectral optimization and the development of targeted multispectral phenotyping which is cheaper and easier to implement for breeding programs on field platforms or UAV systems to enable phenotyping at breeding scale. Together, these approaches can contribute to scalable and generalizable phenomic models capable of supporting grapevine (and other perennial) breeding programs to improve the selection of complex agronomic and quality traits.

## Acknowledgments

The authors are grateful for the support of the lab and field technical assistants from the Department of Plant Breeding during field sampling and phenotyping, as well as support from Anja Rheinberger and Anja Giehl from the Department of Beverage Research at HGU. This work was supported by the LOEWE Professorship for Plant Breeding at Hochschule Geisenheim University funded by the Hessian Ministry of Higher Education, Research and the Arts in Hesse, Germany, awarded to KPVF. Additional support for this research was provided through the “EpiGrape” project funded by the Forschungsring des Deutschen Weinbaus (FDW).

## Data availability statement

All the datasets used to build the predictive models in this study are available as supplementary materials.

## Author contribution

**CARZ:** Conceptualization, Data curation, Formal analysis, Investigation, Methodology, Supervision, Validation, Visualization, Writing – Original draft, Writing – Review and editing

**TS:** Conceptualization, Data curation, Investigation, Methodology, Supervision, Writing – Review and editing

**MS:** Data curation, Writing – Review and editing

**PC:** Data curation, Writing – Review and editing

**HR:** Methodology, Data curation, Writing – Review and editing

**AV:** Data curation, Writing – Review and editing

**KPVF:** Funding acquisition, Project administration, Resources, Writing – Review and editing

## Notes

### Competing Interest Statement

The authors have declared no competing interest.

